# Neurophysiological Measures of Auditory Sensory Processing are Associated with Adaptive Behavior in Children with Autism Spectrum Disorder

**DOI:** 10.1101/2022.02.11.480113

**Authors:** Mairin Cotter, Seydanur Tikir, Ana Alves Francisco, Leona Oakes, Michael J. Crosse, John J. Foxe, Sophie Molholm

**Author notes:** **Corresponding Author** Correspondence to Sophie Molholm.

## Abstract

**Background:** Atypical auditory cortical processing is consistently found in scalp electrophysiological and magnetoencephalographic studies of Autism Spectrum Disorder (ASD), and may provide a marker of neuropathological brain development. However, the relationship between atypical cortical processing of auditory information and adaptive behavior in ASD is not yet well understood.

**Methods:** We sought to test the hypothesis that early auditory processing in ASD is related to everyday adaptive behavior through the examination of auditory event-related potentials (AEPs) in response to simple tones and Vineland Adaptive Behavior Scales in a large cohort of children with ASD (N=89), aged 6-17, and in age- and IQ-matched neurotypically (NT) developing controls (N=120).

**Results:** Statistical analyses revealed significant group differences in early AEPs over temporal scalp regions. Whereas the expected rightward lateralization of the AEP to tonal stimuli occurred in both groups, lateralization of the AEP was only significantly associated with adaptive functioning, in the domains of communication and daily living, in the ASD group.

**Conclusions:** These results lend support to the hypothesis that atypical processing of sensory information is related to everyday adaptive behavior in autism.

## Background

Cortical sensory processing differences in Autism Spectrum Disorder (ASD) may be indicative of aberrant neurodevelopment, and are likely to have cascading effects on higher order cognitive processes (Lewis et al., 2017) that in turn impact clinical phenotype. Studies using electrophysiological (EEG) recordings to examine the brain response to auditory stimulation in ASD consistently reveal smaller and/or slightly delayed auditory evoked potentials (AEP for EEG recordings; auditory evoked magnetic fields (AEMF) for magnetoencephalographic (MEG) recordings) 100-200 ms post stimulus onset over frontal and lateral temporal scalp regions in comparison to age-matched neurotypical (NT) controls (Brandwein et al., 2013; Bruneau et al., 2003; Jansson-Verkasalo et al., 2003; Orekhova et al., 2009; Stroganova et al., 2013). As such AEPs present a strong candidate for a neural marker of cognitive, clinical, and behavioral sequelae of ASD.

Prior work has been directed at exploring relationships between atypical AEPs and the autism phenotype (Brandwein et al., 2015; Foss-Feig et al., 2018), yet very few studies have focused on the relationship between cortical auditory sensory processing and how well a child with a diagnosis of ASD is able to navigate age-appropriate everyday situations (“adaptive behavior”). The Vineland Adaptive Behavior Scales (Vineland) provide an age appropriate measurement of adaptive behavior in the areas of socialization, communication, daily living, motor skills, and maladaptive behavior (Sparrow et al., 2005), and can be used to represent the impact of a neurodevelopmental condition on daily living (Carter et al., 1998). Focusing on the communication domain, Roberts and colleagues (Matsuzaki, Ku, et al., 2019; Roberts et al., 2019) found that the latency of the early auditory MEG response to tonal stimuli was correlated with Vineland adaptive communication scores in a sample of ASD and NT children. Here, we sought to further explore the relationship between auditory processing in ASD and adaptive behavior, by evaluating the relationship between the Vineland domains of socialization and daily living skills in addition to the domain of communication in a large sample of children and adolescents with ASD, using high-density EEG to index auditory sensory processing.

Brain activity in response to tonal and musical stimuli is typically stronger in the right compared to the left cortical hemisphere (E. Orekhova et al., 2013; Parviainen et al., 2019; Yamazaki et al., 2018)) (Matsuzaki, Ku, et al., 2019; Roberts et al., 2019), whereas this pattern is reversed in response to speech and language stimuli (Hornickel et al., 2009; Koyama et al., 2000; Narain et al., 2003). Lateralization of cortical function is observed in many functional domains in humans (Güntürkün et al., 2020; Samara & Tsangaris, 2011) (Dubois et al., 2009; Habas et al., 2012) (Fair et al., 2007), and is often reduced or altered in neurodevelopmental and neuropsychiatric conditions (Berretz et al., 2020; Bishop, 2013; De Guibert et al., 2011; Groen et al., 2008; Qi et al., 2019; Ribolsi et al., 2009; Wexler, 1980). Furthermore, differences in cortical network asymmetries are seen in infants at risk for ASD (Rolison et al., 2021) as well as in sensory processing regions in infants that later go on to receive a diagnosis of ASD (Lewis et al., 2017), and there is extensive evidence for reduced lateralization of language/speech processing in ASD (Floris et al., 2020; Lindell & Hudry, 2013). Studies similarly suggest diminished rightward lateralization for non-speech stimuli in ASD, although to date this has not been extensively reported on (Edgar et al., 2015; Gage et al., 2003; Jorgensen et al., 2021; Matsuzaki, Kuschner, et al., 2019; E. Orekhova et al., 2013; Roberts et al., 2019; Schmidt et al., 2009; Williams et al., 2020). The relationship between auditory lateralization of brain responses to tones and adaptive behavior, however, has not been previously considered.

Here we examined AEPs to simple tones in a cohort of 89 ASD and 120 control participants, ranging in age from 6 to 17 and considered how these responses were related to adaptive behavior. We expected diminished amplitude AEPs in the ASD group compared to the NT group, as well as an atypically lateralized response. Furthermore, we expected that differences in these measures in the ASD group would be associated with poorer adaptive behavior.

## Methods

### Participants

The data presented here were collected at the City College of New York and the Albert Einstein College of Medicine over a 10-year period from 2008–2018. Analyses of subsets of the collected dataset have yielded several publications to date (Brandwein et al., 2013, 2015; Crosse et al., 2019; Cuppini et al., 2020). The sample consisted of children and adolescents with ASD (all were verbal) aged 6–17 and a neurotypically (NT) developing sample matched on age and performance IQ (PIQ). This yielded a sample of 114 participants with ASD (93 males, 21 females) and 142 NT participants (65 males, 77 females). After participants were excluded due to noisy EEG data, poor performance, or too few trials (detailed in Auditory ERP Analysis below), the final sample was 89 ASD participants (72 males, 17 females) and 120 NT participants (55 males, 65 females). Participants were recruited through the Human Clinical Phenotyping Core of the Rose F. Kennedy Intellectual and Developmental Disabilities Research Center, clinician referrals, advertising, and community health fairs. Exclusion criteria included a Performance IQ (PIQ) < 75, abnormal hearing or uncorrected vision, and presence of a neurological disorder. Participants in the NT group were also excluded if they had a neurodevelopmental or neuropsychiatric disorder (as assessed by extensive screening) or had a biological first degree relative with a developmental disorder. Inclusion in the clinical group required an ASD diagnosis confirmed by a trained psychologist, using the Autism Diagnostic Observation Schedule, Second Edition (ADOS-2) (Lord et al., 2012), the Autism Diagnostic Interview-Revised (ADI-R) parent interview, and clinical judgment. In studies that were conducted before 2012, the first edition of the ADOS was used. Intellectual functioning was measured by the Weschler Abbreviated Scale of Intelligence, Second Edition (WASI-II) (Wechsler, 2011). The WASI-II was not administered to 3 ASD and 1 NT participant included in the study. Participants were screened for normal hearing using audiometric threshold evaluation (below 25 dB HL for 500, 100, 2000, 4000 Hz) performed on both ears using a Beltone Audiometer (Model 112). Before beginning the study, parents/ legal guardians gave informed written consent, and participants gave verbal or written assent. The Institutional Review Boards of the Albert Einstein College of Medicine, the City College of New York, and the Graduate Center of the City University of New York approved all procedures and were in accord with the ethical standards as stated in the Declaration of Helsinki.

### Clinical Measures

Adaptive behavior was measured by the Vineland Adaptive Behavior Scale, Second-Edition parent-report questionnaire, which is an assessment tool that measures adaptive behavior for all ages in the domains of socialization, daily living, communication, motor skills, and maladaptive behavior and is an accepted measure of reported adaptive behavior in ASD (Carter et al., 1998; Perry et al., 2009; Ray-Subramanian et al., 2011; Sparrow et al., 2005). Furthermore, the Vineland is applicable to neurotypically developing children, thus allowing us to determine if adaptive behaviors are correlated with measures of auditory neural processing in both groups. In this study, the socialization, daily living, and communication domains were used for analysis. Motor skills and maladaptive domains were excluded because they were not age-appropriate for all participants (motor skills) and/ or are optional (maladaptive) and were not collected for most participants. We also reported the adaptive behavior composite (ABC) scores from the Vineland, which is a combined score of the socialization, daily living, and communication domains, but did not include this total score in analyses as we wished to examine the specific domains of adaptive behavior.

### Data Collection

Clinical and EEG data were collected over 2 visits. In general, clinical data, including the WASI-II and ADOS, were collected during visit 1 in order to confirm an ASD diagnosis and study eligibility, and EEG recording was conducted during the second visit. During this visit, the participants performed an audiovisual simple reaction time task while continuous EEG was recorded from 70 scalp electrodes with an open pass-band from DC to 103 Hz. There were three stimulus conditions presented in random order with equal probability (auditory alone, visual alone, and audiovisual). The “auditory alone” condition consisted of a 1000-Hz tone 75 dBSPL, 5 ms rise/fall time emitted from one speaker (Hartman Multimedia JBL Duet speaker) for 60ms. The visual only condition was an image of a red circle (3.2 cm diameter) which was displayed on a black background 0.4 cm above central fixation along the vertical meridian on a computer monitor (Dell Ultrasharp 1704FTP) for 60 ms at a viewing distance of 122 cm. The audiovisual condition was comprised of the auditory and visual stimuli at the same time. Stimuli were presented in blocks of 100 trials each, and participants were instructed to press a button on a response pad when they saw the instructed stimuli (circle, tone, or both circle and tone) (Brandwein et al., 2015). The three stimuli were presented randomly with an inter-stimulus interval that varied randomly between 1000–3000 ms. Participants were encouraged to take breaks between blocks to preserve focus and prevent fatigue or restlessness. Participants completed between 9 and 11 blocks (the majority completed 10) of 100 trials each with auditory, visual, and audio-visual stimuli randomly presented in each block. Only auditory-alone trials are considered for the current analyses.

### Auditory ERP Analysis

EEG data were analyzed using MATLAB (MATLAB r2020b, MathWorks, Natick, MA) and custom in-house scripts. A low-pass filter of 45 Hz with a slope of 24 dB/octave, and a high-pass filter of 1.6 Hz (Brandwein et al., 2013, 2015) with a slope of 12 dB/octave were applied to each participant’s continuous EEG. Event related potentials (ERPs) for the auditory-alone condition were created by dividing the EEG into 600 ms epochs, including a 100 ms period prior to stimulus onset (−100 ms to 500 ms). Baseline was defined as −100 to 0 ms relative to stimulus onset. Only trials for which the participant pressed the response button were included. Four participants in the ASD group were removed due to having fewer than 100 auditory trials.

Additional participants (7 ASD, 5 NT) were excluded due to excessive electromuscular activity, measured by channels having an amplitude over ±100 µV. Lastly, to detect outliers, we first calculated the maximum amplitude of Global Field Power (i.e., GFP; the standard deviation of all channels) for each subject. Subjects with maximum GFP values that were more than three standard deviations from the mean were excluded (14 ASD, 17 NT). This three-part rejection procedure excluded 18% of our original total dataset (21.9% of ASD group, 15.5% of NT group), leading to our final sample of 89 (72 males, 17 females) ASD participants and 120 (55 males, 65 females) NT participants. For this sample, average number of auditory trials per participant were 288 in the ASD group and 289 trials in the NT group.

The resulting AEPs were referenced to an average of all electrodes. For each participant, electrophysiological indices of early auditory processing were guided by predetermined latency windows over predetermined scalp regions informed by the literature on AEPs (Leavitt et al., 2007, 2011; Shafer et al., 2015). Time-windows and electrodes of interest confirmed (and adjusted if needed) through inspection of the group-averaged ERPs across the dataset. The N1 of the AEP, which we focus on here, can be parsed into subcomponents with positive and negative deflections peaking between ∼70 and 175 ms and with foci over temporal and frontocentral scalp regions. The temporal responses are referred to as the T-complex and include the Ta, a first positive peak, and the Tb, a subsequent negative going response (Tonnquist-Uhlen et al., 2003; Wolpaw & Penry, 1975). A fronto-centrally focused negativity that peaks at about 100 ms is commonly referred to as the N1b (Näätänen & Picton, 1987). Due to previous literature demonstrating group effects in the early auditory response, we focused on these three responses (Brandwein et al., 2013; Bruneau et al., 2003; Orekhova et al., 2009; Williams et al., 2020). For ease of distinction, we refer to these three responses as Ta, Tb, and N1b. For statistical tests analyzing group differences, AEPs were analyzed by region and hemisphere, through left and right temporal channels at 100–125ms and 150–175ms (Ta and Tb, respectively), and through the left and right channels over frontal-central scalp (for a balanced analysis; the fronto-central response tends to be centrally focused, and these right and left channels together are representative of this fronto-central focal region) at 100–125ms (N1b). See Table 1 for electrodes and latency windows used for analysis. For analyses investigating AEP associations with age, severity, and adaptive behavior, the frontocentral N1b was represented by averaging data from the two fronto-centrally placed electrodes (F1 and F2). Lateralization of the auditory response, which was indexed by taking the difference between the responses over left and right temporal scalp regions, was compared across groups for Ta and Tb.

**Table 1.** Latency windows and electrodes used for a priori analysis.

| Component | Latency Window<br>(ms) | Electrodes |
| --- | --- | --- |
| Left Ta | 100–125 | Tp7 |
| Right Ta | 100–125 | Tp8 |
| Left N1b | 100–125 | F1 |
| Right N1b | 100–125 | F2 |
| N1b | 100–125 | F1,F2 |
| Left Tb | 150–175 | Tp7 |
| Right Tb | 150–175 | Tp8 |

### Statistical Analyses

#### Group differences in auditory responses

Statistical analyses were implemented in R (RCoreTeam, 2014). Separate analyses were conducted for responses occurring at 100–125ms, encompassing both Ta and N1b, and for 150-175ms, reflecting the Tb components. For data from the 100–125ms window, a repeated measures ANCOVA was run to examine the main effects of group (ASD and NT), hemisphere (left and right), and region (frontal and temporal) on amplitude. For analyses of the data from the 150–175ms window, where only temporal responses were considered, a repeated measures ANCOVA with group and hemisphere as dependent variables was performed. Due to previous studies demonstrating changes in AEP amplitude throughout development in response to auditory stimuli (Pang & Taylor, 2000; Ponton et al., 2000), age was added as a covariate in both of the ANCOVAs and included in results to further confirm these previous findings. In both of these models, lateralization was defined as the effect of hemisphere on amplitude. For both ANCOVAs, significant outliers were removed to fulfill ANCOVA assumptions. Since outliers can severely affect normality and homogeneity of variance (and ultimately the interpretation of the model), we identified (using a boxplot function in R) and excluded outliers before we ran the models. This process did not exclude a significant number of data points and did not delete entire subjects. Rather, it identified specific rows in the dataset that included outliers. For the repeated measures ANCOVA for 100–125ms, 13 rows were excluded from the 836 rows included in that dataset, and for the 150–175 ms ANCOVAs, 7 rows were excluded from the 418 total rows.

#### Exploratory group analyses

In the above analyses, the probability of Type-I errors was decreased by only considering responses in predetermined time-frames and electrodes. In order to provide a more complete description of our rich dataset, a secondary exploratory phase of analysis was conducted on the full dataset across 64 channels from 50 ms before stimulus onset to 300 ms post stimulus onset. This snapshot of the data allows us to identify potential group differences that were not identified in our a priori analysis and may serve to inform hypothesis generation for future studies. AEPs from the ASD and NT groups were compared using running two-tailed paired t-tests at p <.05 level. Effects lasting at least 16 consecutive ms were then presented in a statistical cluster plot (SCP). This thresholding criterion reduced the likelihood of a type-I error (Guthrie & Buchwald, 1991; Molholm et al., 2002).

#### Clinical associations

To investigate the relationship between auditory responses and adaptive behavior (Vineland socialization, communication, and daily living scores), linear regression models were run for each AEP outcome. These separate models allowed us to better understand potential relationships between specific AEP components and behavior. Due to the aforementioned literature suggesting maturational changes in the AEP from early childhood into early adulthood, age was entered in the first step of the linear regression models.

## Results

### Descriptive Statistics

See Table 2 for participant characteristics of both ASD and NT subjects. Independent t-tests demonstrated that there were no significant differences in age or PIQ between the ASD and NT groups (t [207] = .14, p =.89) and (t [193] =.13, p =.90), respectively. Due to literature suggesting a greater percentage of non-right-handedness in the ASD population (Rysstad & Pedersen, 2016), and a potential effect of handedness on hemispheric lateralization in response to simple stimuli tasks (Papousek & Schulter, 1999; Schmitz et al., 2019), we examined the frequency of left-handed participants in each group. There was a greater percentage of left-handed participants in the ASD group (14.1%) than in the NT group (7.5%). However, chi-square analysis showed that this difference was not statistically significant, X2 (2, N=192) = 2.90, p =.24. In the interest of representing the composition of our study participants, the demographics presented include variables that are not considered in our analyses. As can be seen, our sample was ethnically and racially diverse, although the largest proportion of participants were White (54% of each group). It is also notable that chi-square analyses did not reveal significant group differences in maternal and paternal level of education or race/ethnicity between groups.

**Table 2.**
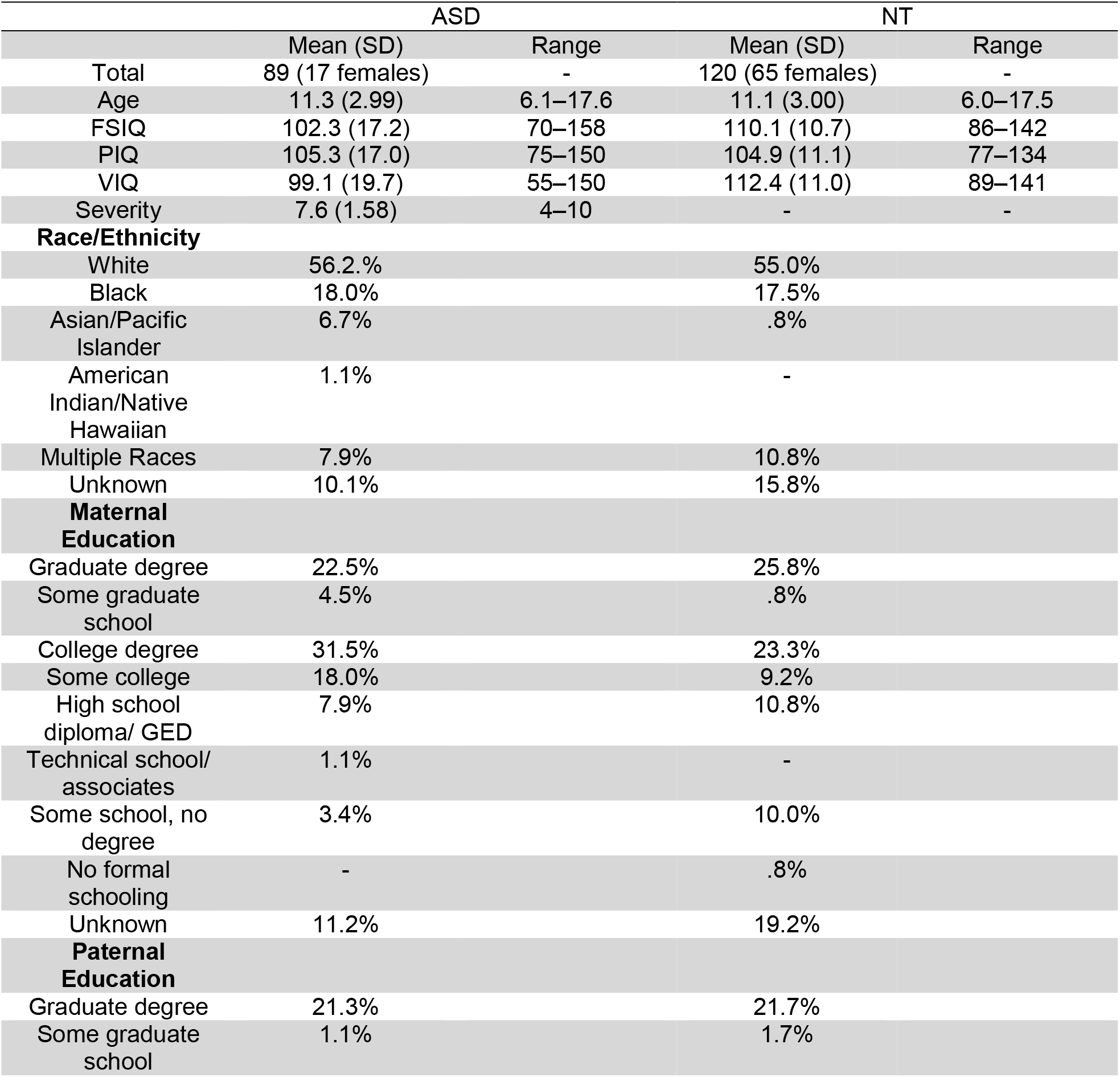
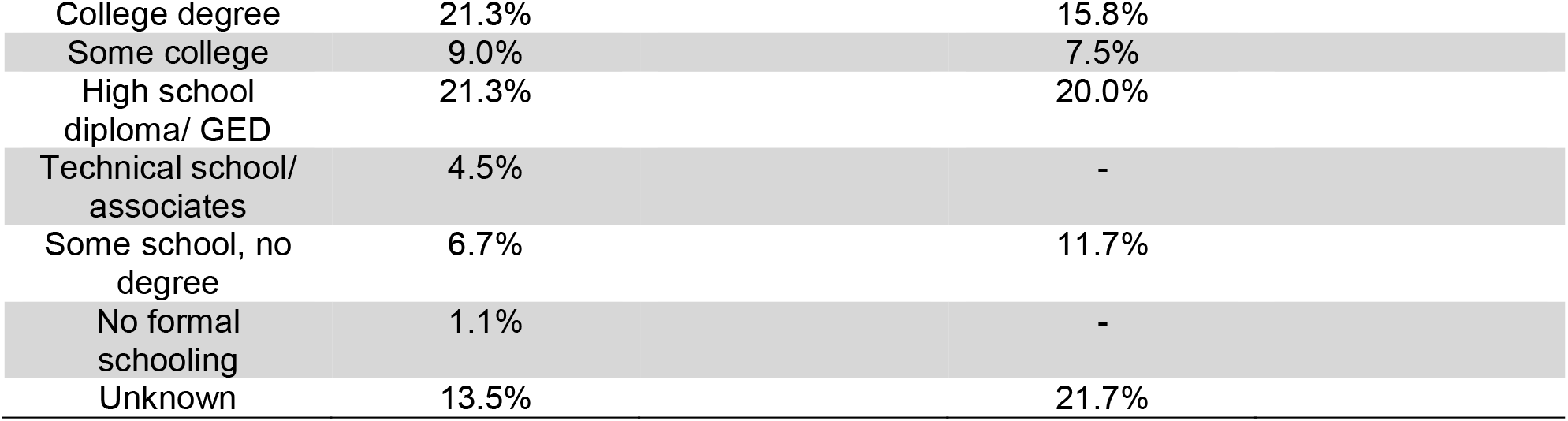
Participant characteristics (mean, range, and standard deviations) after rejection analysis. ASD and NT groups were matched on Performance IQ (PIQ). Verbal IQ (VIQ) and Full-Scale IQ (FSIQ) were significantly different between ASD and NT groups.

While there was a greater percentage of males (80%) in the ASD group compared to the NT group (46%), the male to female ratio in this ASD sample is consistent with asymmetry of males to females diagnosed with ASD in the general population (Loomes et al., 2017). However, this led to an imbalanced sex ratio between the groups. Therefore, clinical and electrophysiological dependent measures were run as a function of sex through independent t-tests and chi-square analyses in the NT group in order to assess whether there was evidence for an influence of sex on the auditory response in the NT group. Sex differences did not attain significance (p > .05) for any of the dependent measures.

### Behavioral Results

Mean hit rates and RTs significantly differed between ASD and NT groups. The ASD group exhibited longer RTs (ASD = 488.98 ms, NT = 435.17 ms), t (207) = −2.76, p =.006, d= .16) and lower hit rates (ASD = 86.35%, NT = 91.39%), t (297) = 3.30, p =.001, d=−.45) compared to the NT group.

### Electrophysiological Results

The morphology of the AEPs appeared to be highly similar between the ASD and NT groups (see Figures 1 and 2), with a positive-going response peaking at ∼100ms (Ta) over bilateral temporal scalp regions, a fronto-central negative-going response peaking at ∼100ms (N1b), and a bilateral temporal negative-going response peaking at ∼150ms (Tb). Nevertheless, visual comparison suggested small group differences in the amplitude of the response over temporal scalp regions. Topographical mapping also suggested rightward lateralization of the response between 75–125ms in both groups, that appeared to be stronger in the NT group.

**Figure 1.**
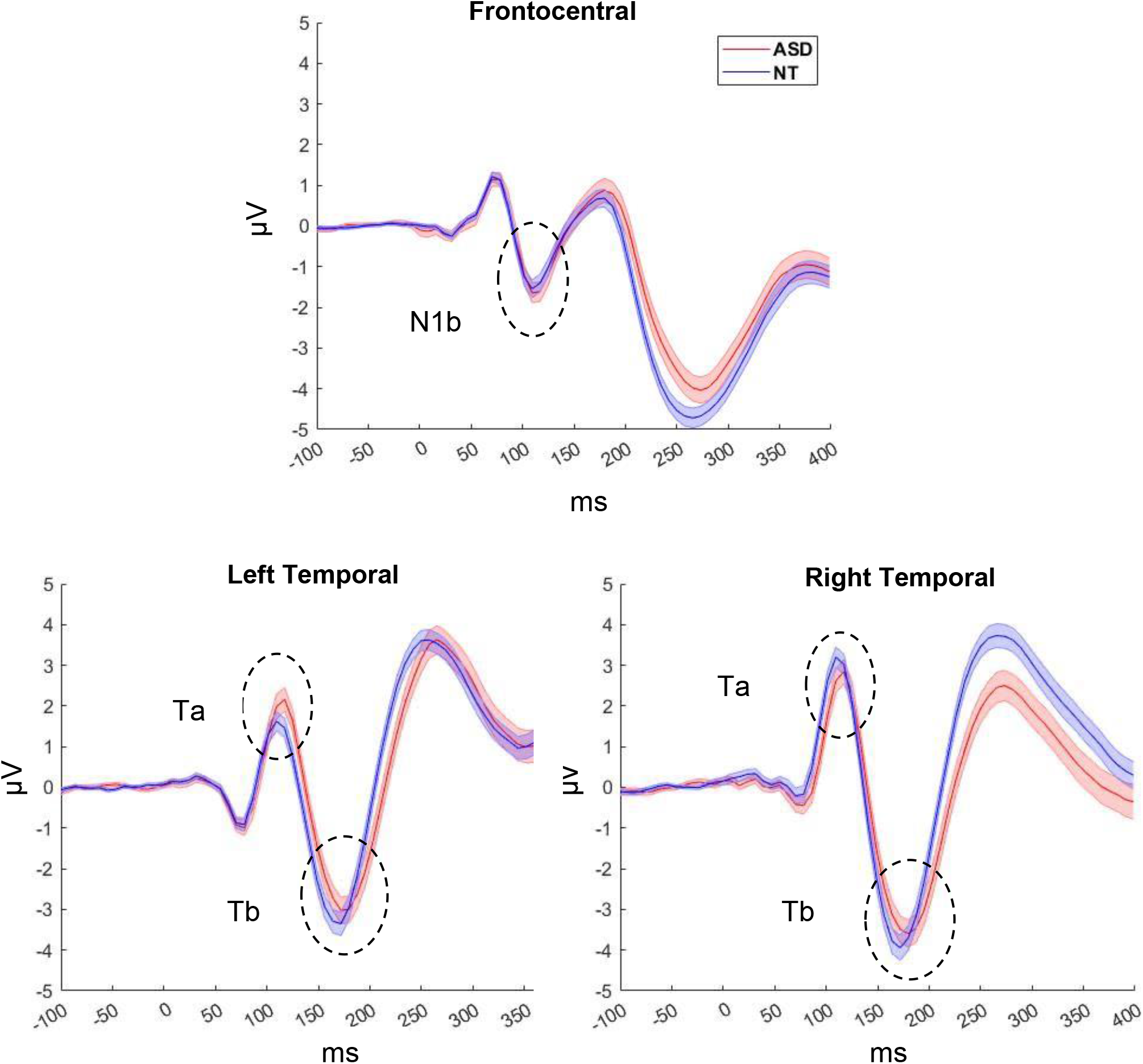
AEP waveforms of mean amplitude for both groups over frontocentral (average of F1 and F2), left temporal (Tp7), and right temporal (Tp8) regions.

**Figure 2.**
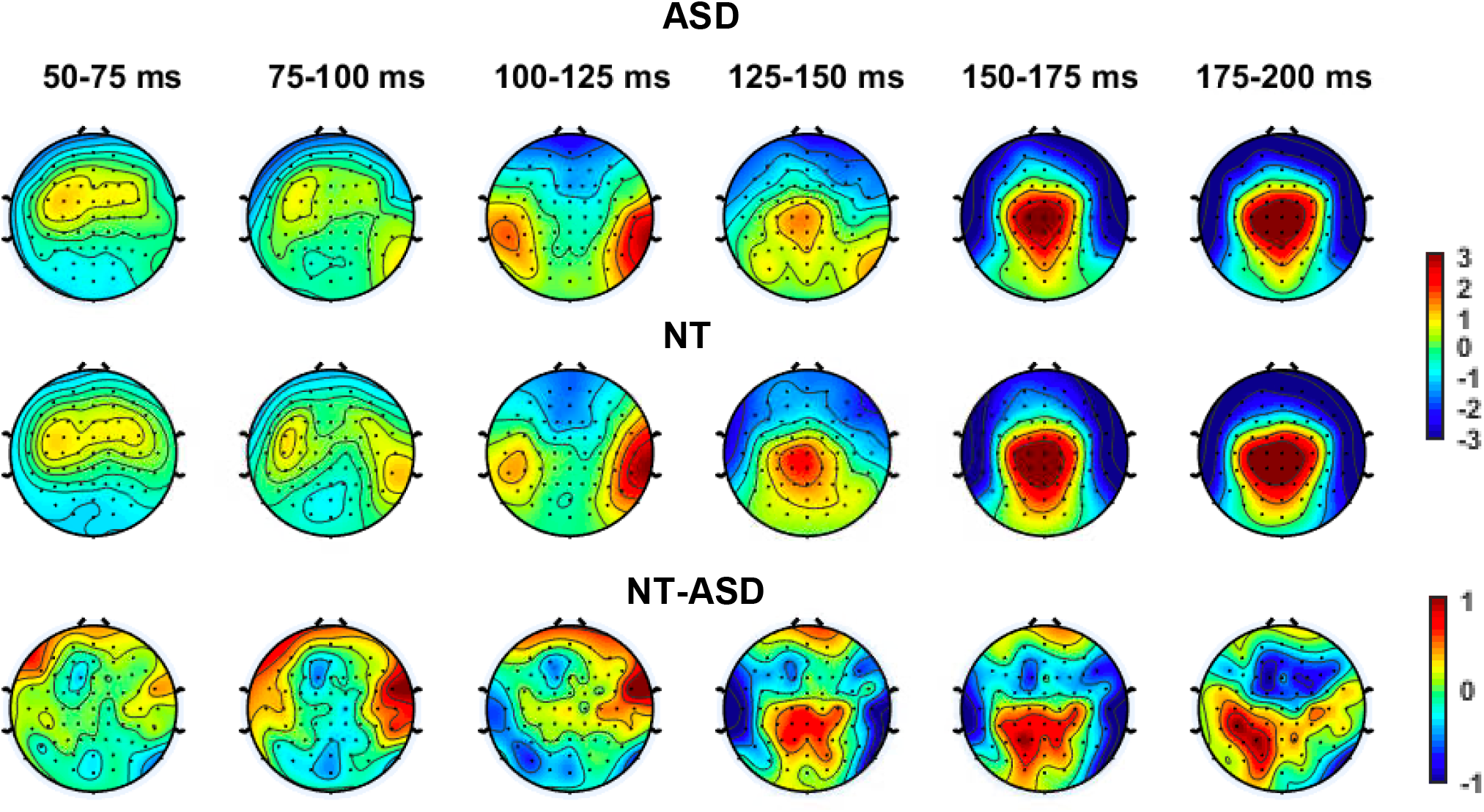
Topographical maps depicting average amplitude of the auditory responses in 25 ms steps from 50–200 ms for ASD, NT, and the difference between ASD and NT. The color bar depicts amplitude in μV.

### Group differences

#### 100–125ms latency window

Levene’s test for equality of variances was significant, and thus we could not assume the data were homogeneous. Therefore, the ANCOVA analysis was adjusted for heteroscedasticity using a coefficient corrected matrix. The ANCOVA revealed significant main effects of age, hemisphere and region (F(1, 807) = 21.92, p <.001, η^2^ =.007, F(1, 807) = 7.99, p =.005, η^2^ =.008, and F(1, 807) = 5663.80, p<.001, η^2^ =.33, respectively). No significant group main effects or interactions were observed. In line with developmental effects on the AEP, there was a significant age × region interaction, F(1,807) = 41.59, p <.001, η^2^ =.032. To further explore this interaction, Spearman correlations were run for the total sample between age and amplitude for each region. Ta and N1b were calculated as an average of the left and right channels. N1b and age were significantly negatively correlated, r_S_=−.35, p <.001 reflecting that the N1b became more negative as age increased. The Ta and age correlation did not reach significance, r_s_ = .089, p =.20. Figure 3 depicts the relationship between age and both N1b and Ta. A second interaction was found for hemisphere and region, F(1, 807) = 16.13, p <.001, η2 =.013. Estimated marginal means demonstrated that the temporal region had the highest difference in amplitude between left and right hemispheres (left = 1.40, right = 2.48) compared to the frontal region (left= −1.08, right = −1.21). This significant hemisphere × region interaction reflects the rightward lateralization of the response over temporal scalp for the total sample. While this was numerically larger in the NT group (see Table 3), this did not reach significance.

**Table 3.** Mean amplitudes of auditory responses over left and right frontal and temporal scalp regions (Ta, N1b, and Tb).

| Window | Region | Hemisphere | ASD Mean (SD) | NT Mean (SD) |
| --- | --- | --- | --- | --- |
| 100–125ms | Frontal (N1b) | Left | −.92 (1.91) | −1.24 (2.22) |
|  |  | Right | −1.17 (1.77) | −1.23 (2.24) |
|  | Temporal (Ta) | Left | 1.52 (2.22) | 1.29 (2.40) |
|  |  | Right | 2.41 (2.15) | 2.56 (2.25) |
| 150–175ms | Temporal (Tb) | Left | −2.14 (2.46) | −2.81 (2.70) |
|  |  | Right | −2.62 (2.51) | −3.40 (2.71) |

**Figure 3.**
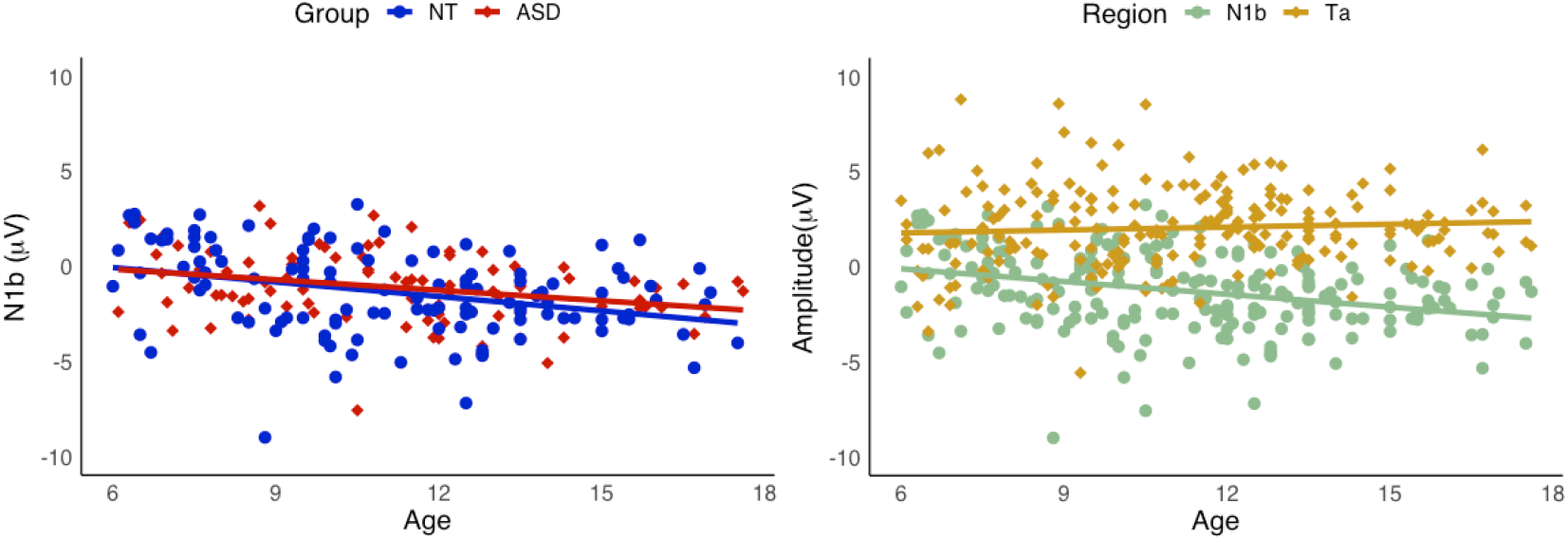
Left panel: Scatterplot of N1b and age by group. Right panel: Scatter plots of N1b and Ta by age group, for total sample. N1b and age were significantly related (rs=−.35, p <.001).

#### 150–175ms latency window

Analysis of responses over temporal scalp regions (Tb) revealed a main effect of group (F(1, 403) = 7.72, p =.006, η^2^ =.018) indicating that the ASD group displayed a decreased (less negative) auditory response over temporal scalp regions compared to the NT group. A main effect of hemisphere was also significant (F(1, 403) = 4.48, p=.03, η^2^ =.011) due to larger responses over right compared to left temporal scalp regions. The group x hemisphere interaction did not reach significance, (F(1, 403) = .056, p=.81, η^2^ =.004). Table 3 displays mean amplitudes by group across hemisphere and region for each time frame analyzed. The ANCOVA did not demonstrate a significant effect for age.

### Exploratory group analyses

Exploratory analysis using the SCP approach suggested early group differences in the 93 to 109 ms window over bilateral temporal/frontal-temporal scalp (channels T7, FT7, FC5; T8, FT8, FC6; see Figure 4), at a time window slightly earlier than the 100–125ms window used in our a priori analysis. Notably, these channels overlapped with the temporal scalp region (and channels) used in our *a priori* analyses. A post hoc ANCOVA was run to confirm this effect with hemisphere and group as independent variables and age as a covariate. A repeated measures ANCOVA was performed on the average amplitude in the 93–109ms window, for averaged data from right and left channels (T8, FT8, FC6 and T7, FT7, FC5 respectively). There were significant group and hemisphere effects, F(1, 398) = 9.01, p =.002, η^2^ =.021, and F(1, 398) = 9.15, p =.003, η^2^ =.022, respectively, and a significant group x hemisphere interaction, F(1, 398) = 4.24, p =.04, η^2^ =.01, due to larger differences between left and right temporal regions in the NT group than the ASD group. See Figures 5 and 6 for plots of significant group differences.

**Figure 4.**
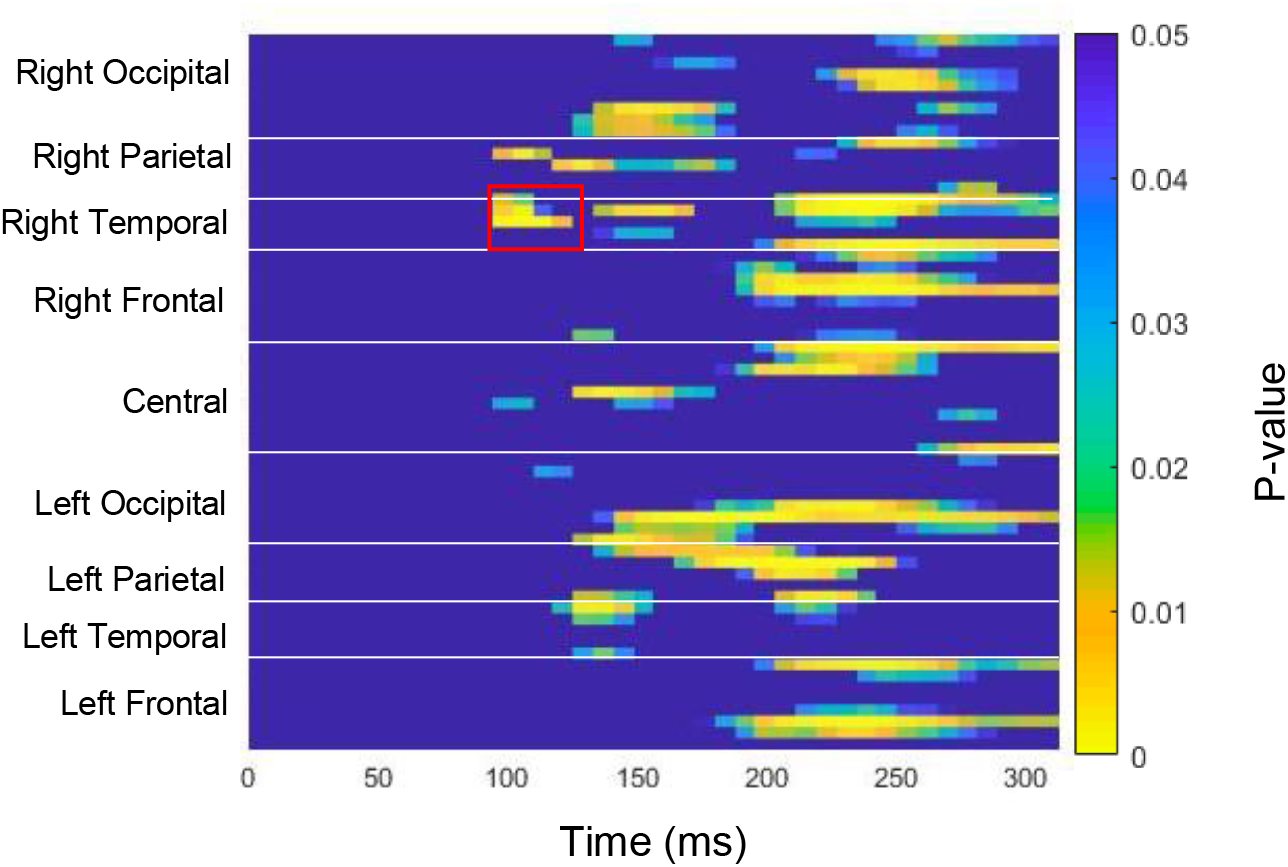
Statistical cluster plots between ASD and NT groups, from 0 to 300ms. The color bar represents significant differences between ASD and NT groups, from p =.05 to p <.001. The red box delineates the 93-109 ms window noted in the text. Channels are grouped by region.

**Figure 5.**
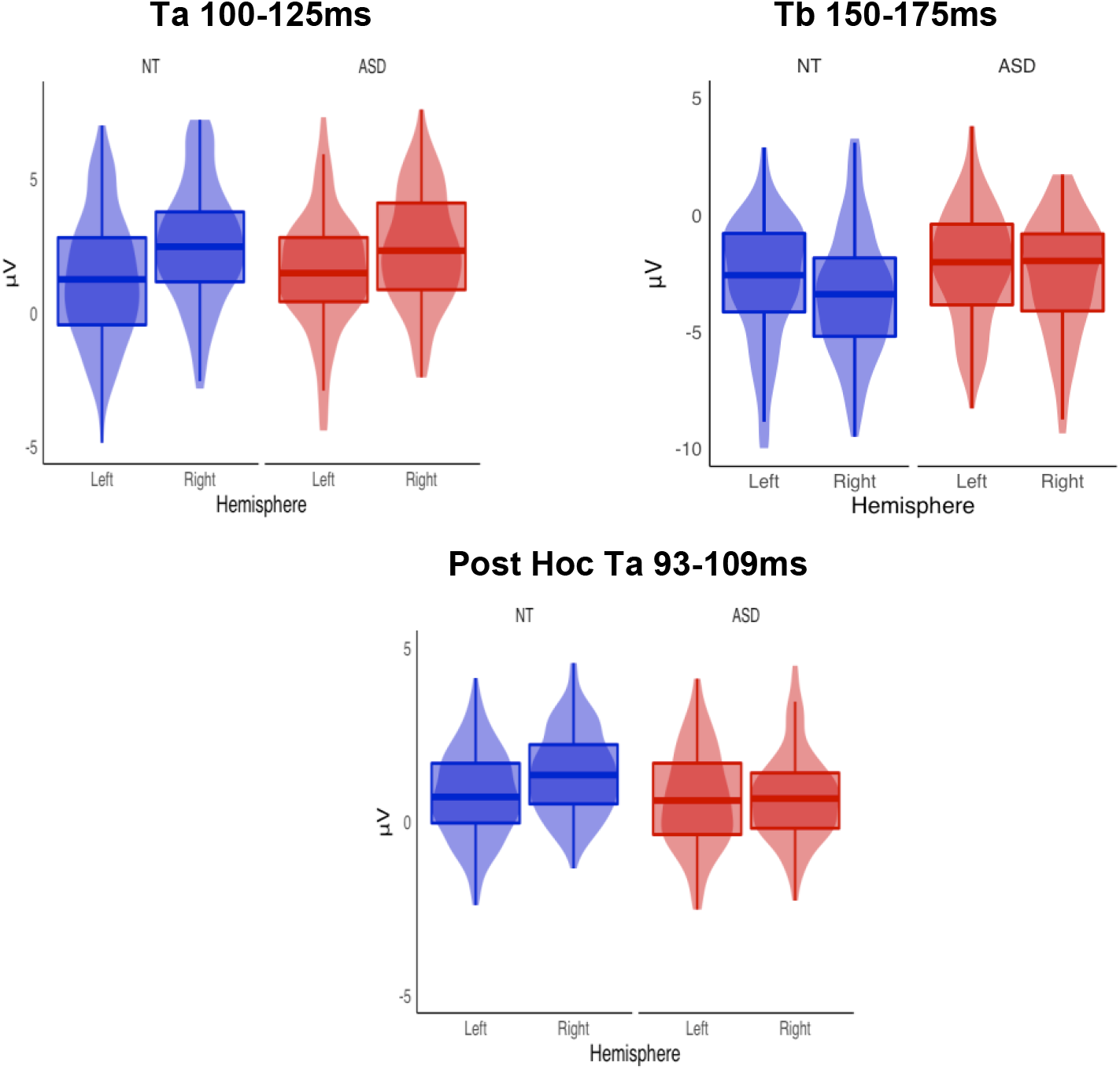
Top panel: Mean amplitudes of Ta and Tb for ASD and NT groups for left and right hemispheres. Bottom panel: Mean amplitudes for ASD and NT groups from post hoc analysis. Scales were adjusted to illustrate distribution.

**Figure 6.**
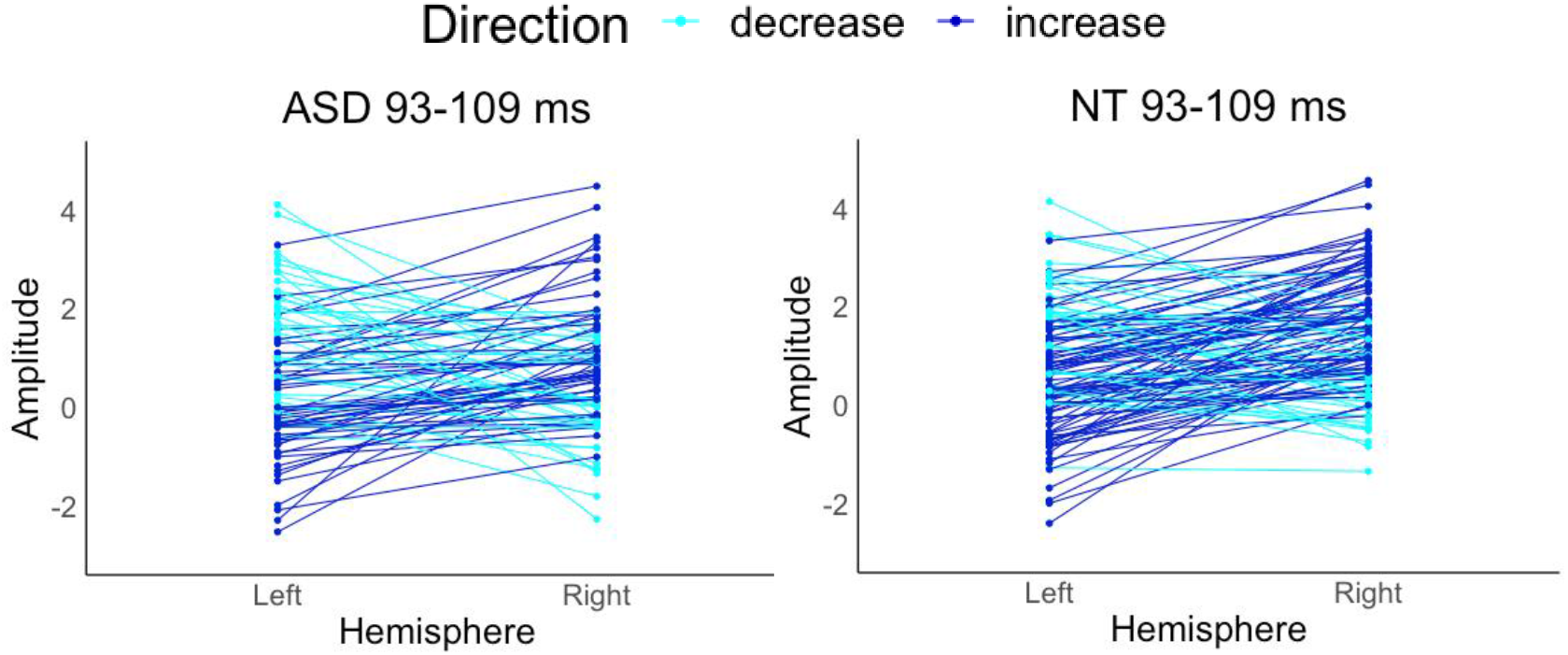
Mean ERPs for each participant across left and right hemispheres in the exploratory post hoc data. The ASD group demonstrated diminished responses in the right hemisphere compared to the NT group.

### Associations between Clinical Measures and Auditory Responses

14 participants in the ASD group and 35 participants in the NT group did not have Vineland scores. Therefore, a subset of this sample consisting of 74 ASD and 44 NT participants, matched on age (range of 6.0–17.5) and IQ (ASD = 102.8; NT = 109.6) was used to investigate the predictive value of measures of adaptive behavior on auditory responses. Chi-square analyses and independent t-tests run between this subset and the participants without Vineland scores did not reveal significant differences on clinical or electrophysiological measures. Within the subset, Vineland scores were significantly lower for the ASD group than the NT group for the communication, socialization, and daily living sub-domains, as well as for the total adaptive behavior composite score, all with p values <.001(see Figure 7).

**Figure 7.**
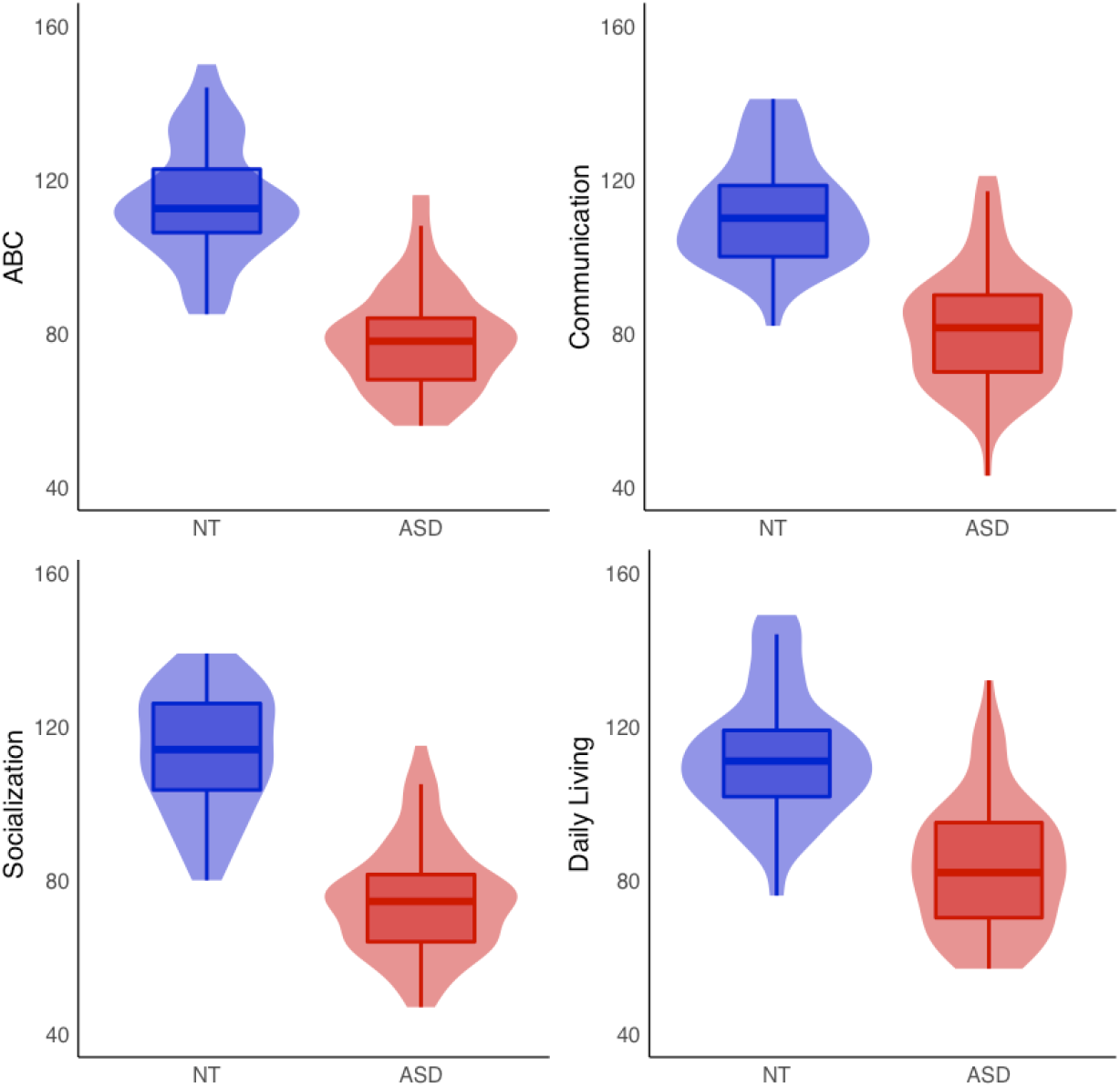
Violin plots of ASD and NT groups for each Vineland domain and total adaptive behavior composite (ABC) score.

Hierarchical linear regression models with age in the first step and clinical measures in the second step were run for each AEP measure across both NT and ASD groups. Models were run for left hemisphere, right hemisphere, and lateralization response for both responses over bilateral temporal regions (Ta and Tb). To ensure that group differences in AEP responses or Vineland measures did not inflate relationships between Vineland domains and AEPs, hierarchical regressions that were significant were conducted separately for ASD and NT groups.

### The Ta Component

For left Ta, Vineland domains of communication, daily living, and socialization were not significant predictors (F(4, 113) = 2.33, p=.06). The total model contributed 7.6% of variance in left Ta amplitude. For right Ta, Vineland domains were not significant predictors (F(4, 113) = 2.32, p=.06). The total model contributed 7.6% of variance in right Ta amplitude. For Ta lateralization, the model was significant, F(4,113)= 5.57, p<.001, contributing 16.5% of variance (p < .001), with Vineland communication and daily living as significant predictors, *B* = −.087, *t*(113) = −4.24, p <.001 and *B* = .051, t(113) = 2.61, p = .01, respectively. Notably, using the data from the exploratory post hoc analysis (93–109ms) yielded similar results, with Vineland communication and daily living significantly predicting Ta lateralization (*B* = −.052, t (113) = −3.22, p =.002 and *B* = .045, t (113) = 2.92, p = .004, respectively.

### The N1b Component

For N1b, the model was significant, F 4, 113) = 2.74, p =.032, with age as a significant predictor, *B* = −.23, *t* (113) = −3.14, *p* = .002, and the Vineland domains contributing only a non-significant additional 1.1% variance (*p* =.22).

### The Tb Component

For left Tb, the total model contributed 4.5% of variance, and the model was not significant for Vineland domains (F(4, 113) = 1.33, *p*=.27). For right Tb, the total model contributed 4.6% of variance, and also did not reach significance for Vineland domains (F(4, 113) = 1.37, *p*=.25). For Tb lateralization, the model was significant, F(4,113) = 2.60, p =.04, contributing 8.4% of variance (*p* =.06), with Vineland communication as a significant predictor, *B* = −.08, *t* (113) = −45, *p* =.009.

### Individual ASD- and NT-group analyses

#### Autism Spectrum Disorder Group

For Ta lateralization in the ASD group, the model was significant after the inclusion of the Vineland domains in the second step, F(4,69)= 3.16, *p*=.019, contributing 15.5% in variance (*p* =.01). In this model, Vineland communication and daily living were significant predictors, *B* = −.093, *t* (69) = −3.18, *p* = .003, and *B* = .007, *t* (69) = 2.99, *p* = .003, respectively. The models were not significant for N1b or Tb lateralization.

#### Neurotypical Group

Models were not significant in the NT group for Ta or Tb lateralization, or N1b. See Figure 8 for residual plots between Ta responses and Vineland domains by group.

**Figure 8.**
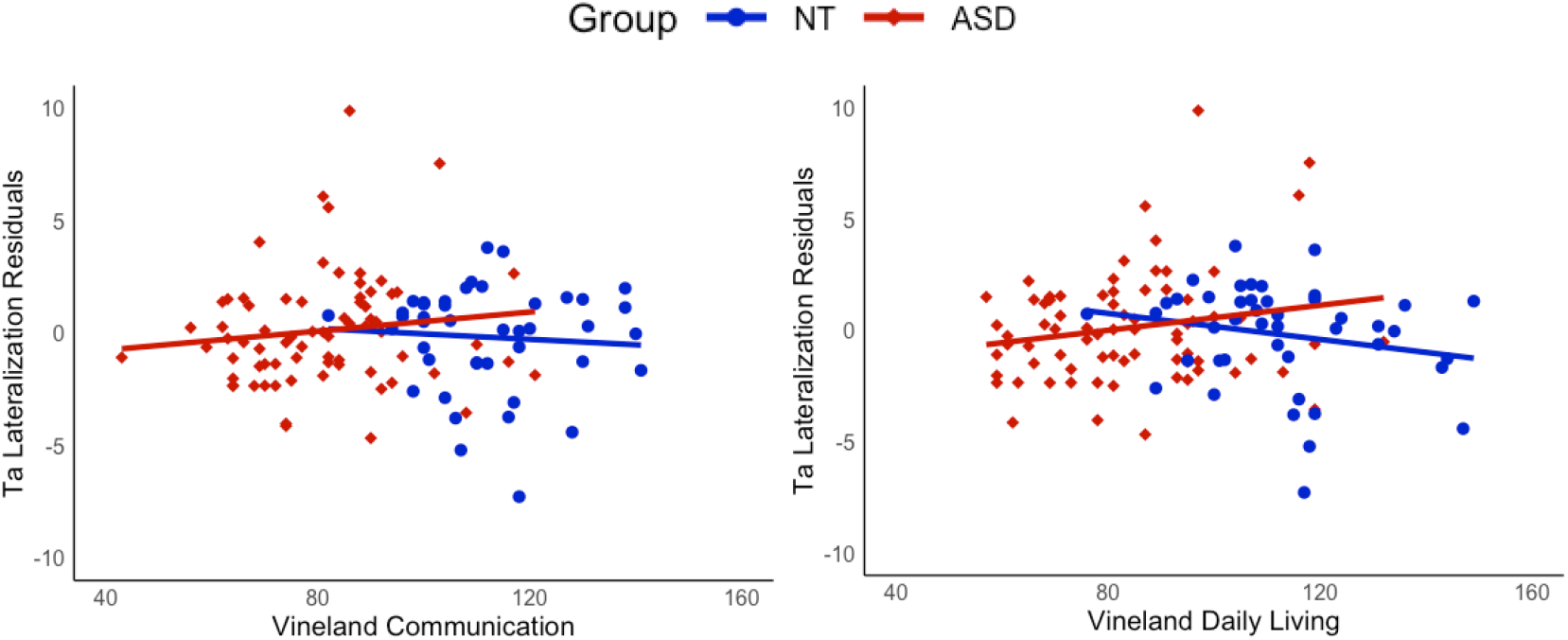
Partial residual plots of Ta amplitude and Vineland communication and daily living.

## Discussion

Engagement in age appropriate adaptive behaviors in everyday situations is significantly reduced in ASD (American Psychiatric Association, 2013). Prior research focusing on electrophysiological responses to auditory stimuli suggests that children and adolescents with ASD exhibit atypical auditory responses (Brandwein et al., 2013; Bruneau et al., 2003; Jansson-Verkasalo et al., 2003; Orekhova et al., 2009; Stroganova et al., 2013). However, how atypical auditory responses are related to adaptive behaviors has not been extensively studied. The current analyses extend our understanding of the relationship between auditory cortical processing and clinical phenotype in ASD by showing that greater rightward lateralization of the early AEP over temporal scalp was associated with better communication and daily living adaptive scores.

With regard to group differences, the ASD group exhibited significantly diminished AEP responses to tones in the 150–175 ms timeframe over bilateral temporal scalp regions. Diminished AEPs align with our previous work (Brandwein et al., 2013) as well as findings from other groups in which responses between 100 and 200ms were of smaller amplitude in ASD compared to control groups (Bruneau et al., 2003; Orekhova et al., 2009b; Williams et al., 2020). While a priori testing of differences for an earlier epoch of 100-125ms failed to reach significance, more comprehensive post hoc consideration of the data supported group differences in a slightly earlier time frame of 93–108. Consistent with previous studies (Brandwein et al., 2013; Gomes et al., 2001; Ponton et al., 2000), age was significantly related to the N1b response in both groups; however this did not interact with group, thus failing to reveal an interaction between childhood development and atypical auditory processing in ASD.

Both groups exhibited the expected rightward lateralization of the AEP to tones. While this lateralization appeared to be reduced in the ASD group (see the 100–125 ms topography maps in Figure 2), this difference did not hold up to statistical testing in our primary analyses. Tellingly, however, the lateralization index derived from the same data revealed a significant relationship with adaptive behavior in the ASD group. This suggests that altered lateralization is an informative feature of ASD, especially when considered at the individual rather than group level. Furthermore, post hoc analysis on a slightly earlier time window supported the presence of altered lateralization in ASD. In prior work from our lab we found atypical lateralization in ASD during a selective attention visual object processing task (Fiebelkorn et al., 2013), and altered lateralization has been observed in language and motor studies (see e.g., (Floris et al., 2016; Leung et al., 2019; Lin et al., 2018; Nickl-Jockschat et al., 2015)). Roberts and colleagues also identified hemisphere specific processing differences in ASD, with the delay in the magnetic auditory evoked response to tones more robust in the right compared to the left auditory cortex in individuals with ASD (Roberts et al., 2019). The finding of altered lateralization across sensory and motor processing and for both language and non-language stimuli suggests a weakening of the hemispheric specialization that is seen in typical development (Floris et al., 2021). As such, this may be a prominent feature of altered neurodevelopment in ASD. Importantly, in line with the idea that greater extent of atypical neural development is likely to be associated with greater clinical severity, our analyses further revealed that greater lateralization of responses to auditory tones was associated with better adaptive functioning in the communication and daily living domains in individuals with ASD.

In addition to the role of aberrant neurodevelopment, how might disruption of auditory processing itself lead to behavioral and perceptual/cognitive sequelae, and in particular delayed development of adaptive behavior? The Vineland daily living domain includes items such as getting dressed or putting away toys. These activities may evoke altered auditory brain responses and specific auditory sensitivities in a child with ASD, preventing them from efficiently completing daily living tasks and developing appropriate adaptive skills. Indeed, hyper- or hypo-reactivity to sensory stimuli is characteristic of ASD (American Psychiatric Association, 2013), and is associated with poorer adaptive behavior (Feldman et al., 2020; Lane et al., 2010; O’Donnell et al., 2012; Rogers et al., 2003). The Vineland communication domain, comprised of subdomains of receptive, expressive and written communication, inherently involves speech and language skills, which are notably impaired in individuals with ASD (Foxe et al., 2015; Lai et al., 2014; Ross et al., 2015). Previous research indicates associations between longer latency MEG responses to auditory tones over the right hemisphere and language ability, as measured by verbal IQ (Matsuzaki, Ku, et al., 2019), as well as Vineland communication skills (Roberts et al., 2019), and between amplitude of the early response and performance during a learned language task in ASD (Arnett et al., 2018). Atypical auditory processing could lead to avoidant reactions to speech, or impair the ability to develop typical communication skills. More research is needed to unpack the respective roles of altered neurodevelopment and atypical auditory processing for adaptive skills.

It is important to note that although a large number of studies investigating early auditory cortical processing in ASD have demonstrated atypical responses in ASD groups compared to NT groups (Brandwein et al., 2013; Bruneau et al., 2003; Jansson-Verkasalo et al., 2003; Matsuzaki, Ku, et al., 2019; E. V. Orekhova et al., 2009; Roberts et al., 2019; Stroganova et al., 2013), these group differences are often modest and the specific results are heterogeneous across studies (Williams et al., 2020), with some studies finding little or no differences (Knight et al., 2020). ASD has a strong genetic basis that is, in most cases, polygenic and variable across individuals (Ramaswami & Geschwind, 2018). Furthermore, unsurprisingly, risk associated allelic variants tend to implicate neurobiological pathways involved in fetal neural development and synaptic function (de la Torre-Ubieta et al., 2016). Consistent with this heterogeneity, prior work suggests that atypical connectivity in sensory and higher order networks in ASD is highly idiosyncratic compared to controls (Benkarim et al., 2021; Hahamy et al., 2015). Thus, while the auditory cortex appears particularly vulnerable to resulting neuropathology, just how this plays out may vary based on an individual’s genetic background, specific set of genetic vulnerabilities, and the environmental factors that they are exposed to.

## Conclusions

This study supports prior findings that children and adolescents with ASD exhibit atypical auditory responses at the neural level. Furthermore, our results support a relationship between atypical auditory processing in children and adolescents with ASD and adaptive behavior. Although there is great heterogeneity in the ASD population, these findings utilizing a large dataset with a wide range of ages indicate the presence of a relationship between basic neuropathological processes and maladaptive behavior in this population. Future studies will be needed to understand how this knowledge can inform approaches to improving adaptive function in this group, especially in the domains of communication and daily living skills. Furthermore, expanding the group to include minimally verbal and nonverbal indiviuals with ASD will be an important step to expanding understanding of how auditory processing differences contribute to adaptive behavior and other aspects of the autism clinical phenotype.

## List of Abbreviations

ADI-R: Autism Diagnostic Interview-Revised
AEMF: Auditory evoked magnetic fields
ASD: Autism spectrum disorder
ADOS-2: Autism Diagnostic Observation Schedule-2
EEG: Electroencephalography
ERP: Event related potential
FSIQ: Full-scale intelligence quotient
MEG: Magnetoencephalography
NT: Neurotypical
PIQ: Performance intelligent quotient
VIQ: Verbal intelligent quotient
WASI-II: Weschler Abbreviated Scale of Intelligence, Second Edition
AEP: Auditory Evoked Potential
SCP: Statistical Cluster Plot
ABC: Adaptive Behavior Composite
RT: Reaction Time

## Declarations

### Ethics approval and consent to participate

All procedures were approved by the Institutional Review Boards of the Albert Einstein College of Medicine, the City College of New York, and the Graduate Center of the City University of New York and were in accord with the ethical standards as stated in the Declaration of Helsinki.

### Consent for publication

Not applicable.

### Availability of data and materials

Data from the findings of this study are available from the authors upon request. The authors will make the Matlab scripts available in a public repository (Github).

### Authors’ contributions

Conceived and designed the study: J.J.F. and S.M. Data analysis: M.C., S.T., A.A.F., and M.J.C. Supervision: S.M. Writing-original manuscript preparation: M.C. and S.M. Writing-review and editing: S.M., A.A.F., S.T., L.O., M.C., M.J.C. and J.J.F. All authors read and approved the final manuscript.

Acknowledgments: We extend our deep appreciation to all of the families that have generously given their time to participate in this research. This work could not be done without the clinicians that have performed or supervised clinical and cognitive testing including: Alice Brandwein; Juliana Bates; Hilary Gomes, and Natalie Russo. We are grateful to the research assistants and technicians that have put in great efforts to collect high quality EEG data while ensuring the comfort of our participants, including: Douwe Horsthuis, Alaina Berruti, Frantzy Acluche, Greg Peters, Sarah Ruberman, and Elise Taverna. An earlier version of this manuscript was submitted to the Department of Psychology at Fordham University towards fulfillment of the first author’s master’s degree.

